# Nonheme Fe Enzyme-Catalyzed Enantioselective and Regiodivergent C(sp^3^)–H Amidation: Biocatalytic Asymmetric Synthesis of 1,2- and 1,3-Diamine Derivatives

**DOI:** 10.64898/2026.11.17.767434

**Authors:** Hongda Chen, Shijia Li, Huichong Liu, Liu-Peng Zhao, Ken Lin, Peng Liu, Yang Yang

## Abstract

Nonheme Fe enzyme-catalyzed asymmetric, site-selective C–H amidation remains underexplored. Here we report the repurposing and directed evolution of the plant-derived nonheme Fe enzyme 1-aminocyclopropane-1-carboxylic acid oxidase (ACCO) into a family of biocatalysts for highly regio- and enantioselective C(sp^3^)– H amidation. Engineered ACCO variants catalyzed the asymmetric synthesis of cyclic ureas from *N*-(benzoyloxy)urea substrates, which are readily prepared from aliphatic amines, through the activation of benzylic, allylic, propargylic, and unactivated aliphatic C(sp^3^)–H bonds, including enantioconvergent amidation of racemic tertiary C(sp^3^)–H substrates. Directed evolution further generated complementary enzymes that selectively promoted either 1,5- or 1,6-hydrogen atom transfer, enabling catalyst-controlled, regiodivergent enantioselective synthesis of both 1,2- and 1,3-diamine derivatives from common precursors. The evolved biocatalysts exhibited total turnover numbers of up to 4,500 and were readily applicable to gram-scale whole-cell biotransformations. Mechanistic investigations supported a stepwise mechanism involving a carbon-centered radical intermediate. Kinetic isotope effect and isotopically labeled substrate studies indicated C(sp^3^)–H bond cleavage as the rate-limiting step, while stereochemical analyses revealed that the overall enantioselectivity is determined by the C–N bond-forming radical rebound rather than hydrogen atom transfer. These results established nonheme Fe enzymes as a versatile platform for selective C–H amidation and further expanded the scope of new-to-nature nonheme Fe biocatalysis for asymmetric C–H functionalization.

**Entry for the Table of Contents:** An enantioselective and regiodivergent biocatalytic C(sp^3^)–H amidation of urea derivatives is developed. Engineered variants of the nonheme iron enzyme ACCO control 1,5-versus 1,6-hydrogen atom transfer, affording chiral five- and six-membered cyclic ureas and providing access to valuable 1,2- and 1,3-diamine derivatives.

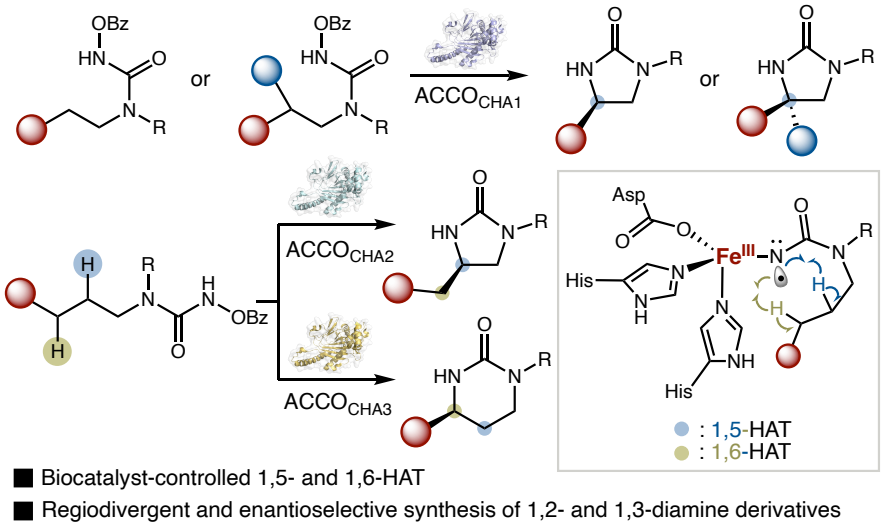

## Introduction

Catalytic regio- and stereoselective C–H functionalization represents a key frontier in modern synthetic organic chemistry^1^ and biocatalysis.^2^ By enabling catalyst-controlled, site- and stereoselective transformation of otherwise inert C–H bonds, these technologies hold the potential to streamline the synthesis of valuable, stereochemically defined molecules from readily available precursors. Over the past decade, important studies by Arnold, Fasan, Hartwig and other researchers established biocatalytic nitrene transfer as a general strategy for asymmetric C–H amination.^3,4^ To date, a range of biocatalytic intramolecular and intermolecular C(sp^3^)–H amination reactions has been developed to convert various hydrocarbon and functionalized substrates into useful aminated products.^3,4^ For a long time, heme proteins,^5^ including cytochromes P450, globins and cytochromes *c*, have been regarded as the major workhorse biocatalysts for asymmetric C–H amination.^4^ In contrast, the development of nonheme Fe enzymes^6^ for regio- and stereoselective C–H amination reactions has lagged behind,^7^ in part due to the less well-defined coordination environments of the Fe center in these nonheme systems compared with those in extensively studied heme enzymes. Furthermore, the mechanism and origin of enantioselectivity in these nonheme Fe enzyme-catalyzed new-to-nature C–H amination reactions remain to be further elucidated. Thus, further investigation and development of nonheme Fe biocatalysts may provide a valuable complement to heme enzymes^4^ as well as well-tailored small-molecule transition-metal catalysts^8,9^ for selective C–H amination.

Our laboratory has been investigating the repurposing and engineering of nonheme Fe enzymes for asymmetric C–H functionalization reactions not previously known in native enzymology.^10,11^ In 2024, our group and the Huang group reported the repurposing and directed evolution of nonheme Fe enzymes to catalyze asymmetric C(sp^3^)–H fluorination via a fluorine atom transfer mechanism.^10^ Subsequently, inspired by elegant studies from the Meggers laboratory on transition-metal-catalyzed nitrogen migration reactions,^8^ our group and other researchers engineered a suite of nonheme Fe biocatalysts to facilitate nitrogen migration reactions for the asymmetric synthesis of *N*-protected amino acids.^11^ Based on these results, we envisioned leveraging the nonheme Fe enzyme platform to enable a broader range of synthetically valuable enantioselective C–H functionalization reactions, particularly those involving challenging site selectivities eluding the state-of-the-art chiral small-molecule Fe catalysts.

In this context, inspired by transition-metal-catalyzed C–H amination,^8,9^ we recently questioned whether we could repurpose and evolve nonheme Fe enzymes to catalyze the regio- and enantioselective C–H amidation of aliphatic amine derivatives, thereby allowing the formal asymmetric and divergent synthesis of 1,2- and 1,3-diamine derivatives with widespread applications in the pharmaceutical and agrochemical industries as well as organic synthesis (Figure 1). In this strategy, appending an *N*-(benzoyloxy)formyl directing group to widely available aliphatic amines would afford *N*-(benzoyloxy)urea substrates.^9e,f,g^ Reaction of the *N*-(benzoyloxy)urea substrate with the nonheme Fe center via α-elimination would afford the key Fe nitrenoid intermediate with substantial spin density on the nitrogen center. Subsequent hydrogen atom transfer (HAT) would convert this Fe nitrenoid species into a carbon-centered radical and an Fe–NHR intermediate. Finally, radical rebound involving this nascent carbon-centered radical would furnish the enantioenriched cyclic urea product, which is readily transformed into a chiral diamine using established procedures.^9e,f,g^ Overall, this traceless directing group strategy enables the formal conversion of simple aliphatic amines into valuable chiral diamines. Beyond the utility of these chiral diamines, chiral cyclic ureas are also ubiquitous structural elements in bioactive small-molecule compounds (Figure 1(A)).

**Figure 1.**
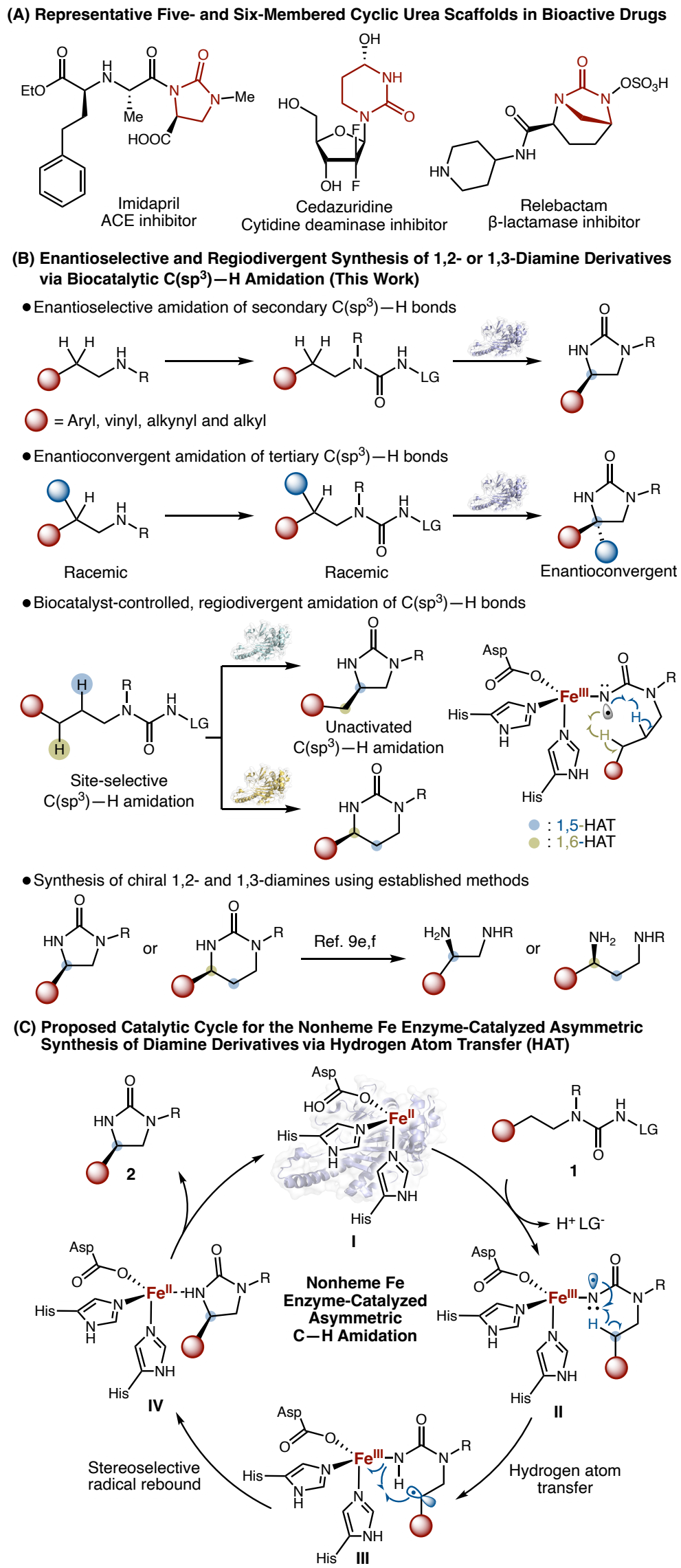
Nonheme Fe Enzyme-Catalyzed, Regiodivergent and Enantiose-lective C(sp^3^)–H Amidation: Asymmetric Synthesis of 1,2- and 1,3-Diamine Derivatives. (A) Selected examples of five- and six-membered cyclic ureas in bioactive drugs. (B) Nonheme Fe enzyme-catalyzed enantioselective and regiodivergent synthesis of 1,2- or 1,3-diamine derivatives via biocatalytic C(sp^3^)–H amidation. (C) Proposed catalytic cycle for the nonheme Fe enzyme-catalyzed asymmetric synthesis of diamine derivatives.

Herein, we show that nonheme Fe enzymes could be repurposed and evolved to catalyze the asymmetric synthesis of diamines through enantioselective amidation of a variety of C(sp^3^)–H bonds, including secondary benzylic, allylic, propargylic, and unactivated aliphatic C(sp^3^)–H bonds. Furthermore, racemic substrates bearing a tertiary C(sp^3^)–H bond could also be transformed into the corresponding α-tetrasubstituted aminated products in an enantioconvergent fashion. In particular, we sought to engineer an orthogonal set of regiodivergent nonheme Fe enzymes to achieve the enantioselective synthesis of both 1,2- and 1,3-diamines via an enzyme-controlled 1,5- and 1,6-HAT mechanism, thereby expanding the growing repertoire of enzyme-controlled regiodivergent C–H amination.^4b,11e^ Such catalyst-controlled, site-selective asymmetric C(sp^3^)–H amidation reactions continue to present a significant challenge for the state-of-the-art small-molecule transition-metal catalysts. Overall, the development of general, efficient, and regio and enantioselective nonheme Fe biocatalysts highlighted their synthetic potential for accomplishing synthetically useful selective C–H functionalization. Furthermore, these newly engineered nonheme Fe biocatalysts contribute to the growing catalytic repertoire for transition-metal-catalyzed C(sp^3^)–H amidation, providing a biocatalytic alternative for efficient and enantioselective amidation (see Table S18 for a comparison of the state-of-the-art methods).

## Results and Discussion

We commence our study by evaluating our in-house collection of nonheme Fe enzymes for the intramolecular C(sp^3^)–H amidation of benzoyloxyurea **1a**, a transformation proposed to proceed via a 1,5-hydrogen atom transfer (1,5-HAT) step (Figure 2(A)). Among the nonheme enzymes evaluated, hercynine oxygenase from *Mycolicibacterium thermoresistibile* (*Mth*EgtB, UniProt ID: G7CFI3),^12^ which features a three-histidine facial triad, furnished the desired 2-imidazolidinone product **2a** in 30% yield with a 31:69 enantiomeric ratio (e.r.) favoring the (*S*)-enantiomer (Figure 2(A), entry 1). Catechol 2,3-dioxygenase from *Rhodococcus aromaticivorans* DK17 (C23O, UniProt ID: Q6REQ5),^13^ which features a 2-His-1-carboxylate facial triad, afforded **2a** in 3% yield and 51:49 e.r. (entry 2). Isopenicillin N synthase from *Emericella nidulans* (*Eni*IPNS, UniProt ID: P05326),^14^ which also contains a 2-His-1-carboxylate facial triad, afforded **2a** in 6% yield and 47:53 e.r. (entry 3). Metapyrocatechase from *Pseudomonas putida* mt-2 (MPC, UniProt ID: P06622)^15^ furnished racemic **2a** in 3% yield (50:50 e.r., entry 4). Deacetoxycephalosporin C synthase from *Streptomyces clavuligerus* (*Sc*DAOCS, UniProt ID: P18548)^16^ afforded **2a** in 10% yield and 25:75 e.r. (entry 5). 1-Aminocyclopropane-1-carboxylic acid oxidase from *Petunia hybrida* (*Phy*ACCO, UniProt ID: Q08506)^17^ also catalyzed the formation of **2a** in 30% yield, albeit with a low level of enantioselectivity (47:53 e.r., entry 6). Collectively, these results demonstrated the utility of a range of nonheme Fe enzymes with diverse native biological functions toward this non-native asymmetric intramolecular C(sp^3^)–H amidation (see Table S3 for further details). Among all the nonheme Fe enzymes we examined, our previously evolved *Phy*ACCO variant ACCO_Nim1_ containing eight additional mutations relative to wt *Phy*ACCO, (*Phy*ACCO G156E K158T K172V G173R R175P I184A L186V V236G), which was engineered for the enantioselective synthesis of noncanonical α-amino acids,^11a^ exhibited substantially enhanced catalytic activity, affording **2a** in 80% yield and 91:9 e.r. in favor of the opposite (*R*)-enantiomer. The related ACCO_Nim2_^11a^ (*Phy*ACCO G156E K158T K172V G173V R175P I184A L186V V236K F250L) provided a higher yield of 81%, albeit with a slightly reduced enantioselectivity (77:23 e.r.).

**Figure 2.**
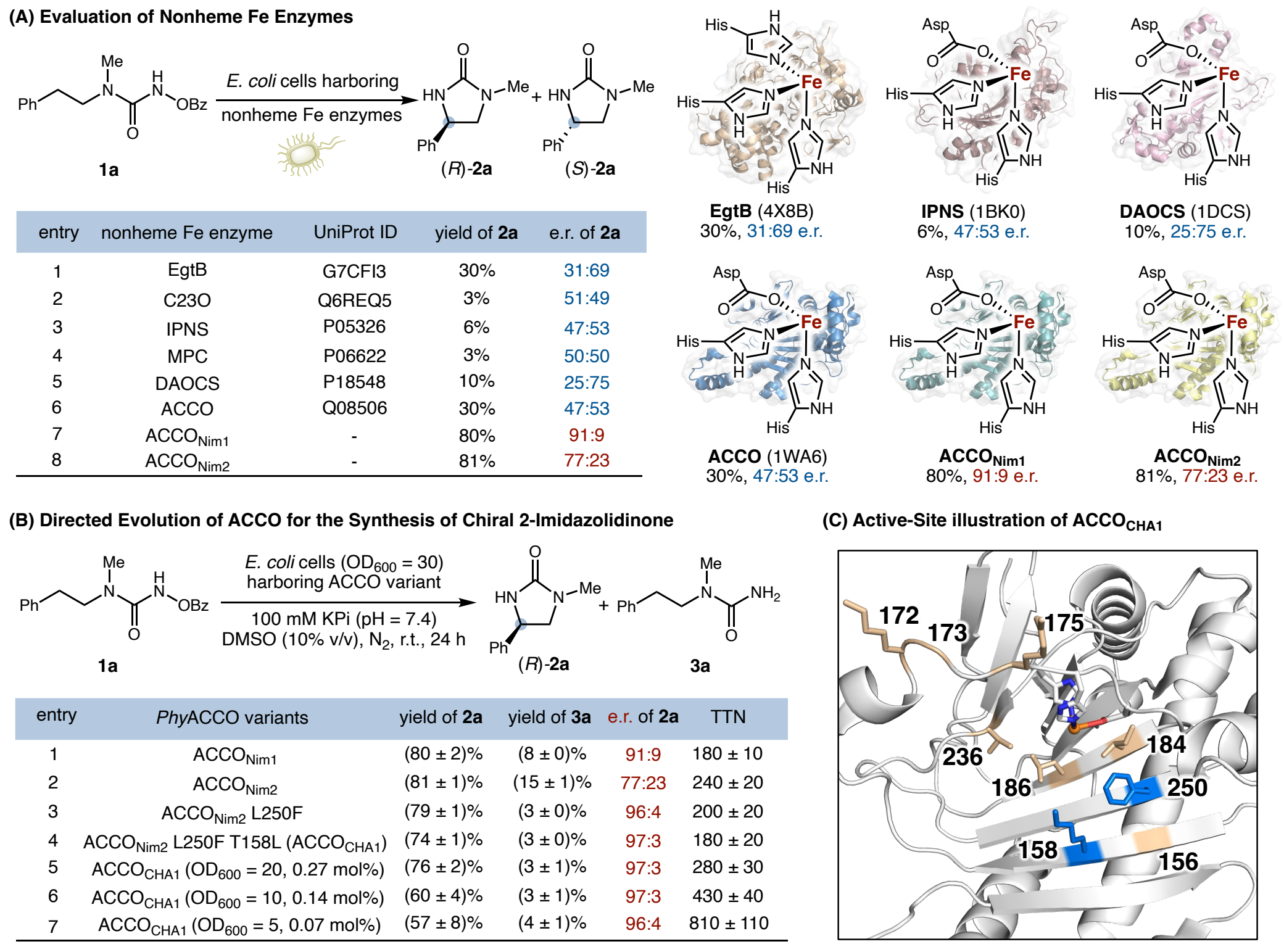
Discovery and Engineering of ACCO_CHA1_ for Enantioselective C(sp^3^)–H amidation. (A) Evaluation of nonheme Fe enzymes for biocatalytic asymmetric C(sp^3^)–H amidation using whole *E. coli* cells. (B) Protein engineering of ACCO_Nim2_. Reaction conditions: 1a (10.0 mM), *E. coli* cells harboring ACCO variants in KPi buffer (100 mM, pH = 7.4) at the indicated OD_600_, DMSO (10% v/v), room temperature, 24 h. All the biocatalytic reactions were performed as technical triplicates. (C) Active-site illustration of ACCO (PDB ID: 1WA6) and beneficial mutations identified from protein engineering. Marine residues denote beneficial mutations identified in ACCO_CHA1_. Wheat residues denote mutations of ACCO_Nim2_ relative to wt ACCO.

Further screening of our in-house collection of variants from our ACCO_Nim_ lineage^11a^ identified ACCO_Nim2_ L250F as the most promising variant, exhibiting excellent activity and highest enantioselectivity among all the nonheme Fe enzymes examined (79% yield and 96:4 e.r., entry 3). We hypothesized that the side chain of L250F resides close to the Fe-bound substrate, thereby playing an important role in controlling enzyme activity and enantioselectivity. With ACCO_Nim2_ L250F as the parent, we initiated a brief enzyme engineering effort using site saturation mutagenesis (SSM) and screening by randomizing residues 158, 173, 186 and 250. Guided by the crystal structure of *Phy*ACCO (PDB ID: 1WA6),^17^ we targeted active-site and substrate tunnel residues proximal to the nonheme Fe center (Figure 2C). Each SSM library was constructed using the 22c-trick method,^18^ and 88 clones from each library were screened. Protein expression and preliminary activity screening were performed in 96-well plates, and promising variants were subsequently validated using bacterial cultures grown in Erlenmeyer flasks. During this round of engineering, we identified a new variant ACCO_Nim2_ L250F T158L, further improving the enantioselectivity of **2a** to 97:3 e.r. (entry 4). The evaluation of other SSM libraries did not lead to a significant improvement (Tables S5–S7). ACCO_Nim2_ L250F T158L was designated ACCO_CHA1_ (ACCO C–H amidase 1). Further reducing the loading of *E. coli* suspension overexpressing ACCO_CHA1_ to an OD_600_ of 5 (ca. 0.07 mol% ACCO_CHA1_), afforded **2a** in 57% yield, 810 ± 110 total turnover number (TTN) and 96:4 e.r. (entry 7), demonstrating the potential utility of ACCO_CHA1_ whole-cell biocatalyst for laboratory-scale synthesis. The absolute stereochemistry of **2a** was confirmed by comparison with literature precedents.^9e^

With the engineered variant ACCO_CHA1_ in hand, we investigated the substrate scope of this biocatalytic asymmetric C(sp^3^)–H amidation for the synthesis of 2-imidazolidinones (Figure 3). It was found that *para*-substituents (**2b**) on the aromatic ring were compatible. Additionally, a range of halogen substituents, including a fluoro-(**2c** and **2g**), a chloro-(**2d**), a bromo-(**2e**), and an iodo-(**2f**) group, were also tolerated, providing useful functional group handles for subsequent cross-coupling diversification.^19^ Substrates possessing aromatic electron-donating substituents such as a *para*-methoxy group (**2h**), and electron-withdrawing substituents including a *para*-cyano (**2i**) and a *para*-trifluoromethyl (**2j**), also furnished the corresponding 2-imidazolidinones in good yields and enantioselectivities. Heterocyclic substrates including a 2-thienyl (**2k**) and 3-thienyl (**2l**) moiety could also be transformed in a highly enantioselective fashion. A sterically demanding naphthyl moiety (**2m**) was also compatible, affording the corresponding imidazolidinone product in excellent yield and enantiomeric ratio. The nonheme Fe biocatalyst was readily applicable to the amidation of allylic (**2n** and **2o**) and propargylic (**2p**) C(sp^3^)–H bonds, affording enantioenriched C(sp^3^)–N bond-forming products with modest to excellent enantioselectivities. No olefin aziridination products were observed in the biocatalytic transformation of **1n** and **1o**, further demonstrating the nonheme enzyme’s excellent chemoselectivity. A slightly larger *N*-ethyl group (**2q**) was also tolerated by the current biocatalytic protocol. 5,5-Fused bicyclic product **2r** could also be prepared enantioselectively using an indanyl substrate. Furthermore, without additional engineering, the amidation of unactivated secondary C(sp^3^)–H bonds (**2s**) could also be achieved with ACCO_CHA1_, albeit with modest activity and enantioselectivity. We expect that further directed evolution could improve the transformation efficiency of this class of substrates. Collectively, these results demonstrated the synthetic utility of ACCO_CHA1_ in the enantioselective amidation of a wide variety of secondary C(sp^3^)–H systems. We note that whole *E. coli* cells promoted the conversion of the benzoyloxyurea substrates (**1**) to the corresponding reduced ureas (**3**) in the absence of ACCO_CHA_ variants in 5–10% yield, resulting in slightly reduced product yields in these whole-cell reactions.

**Figure 3.**
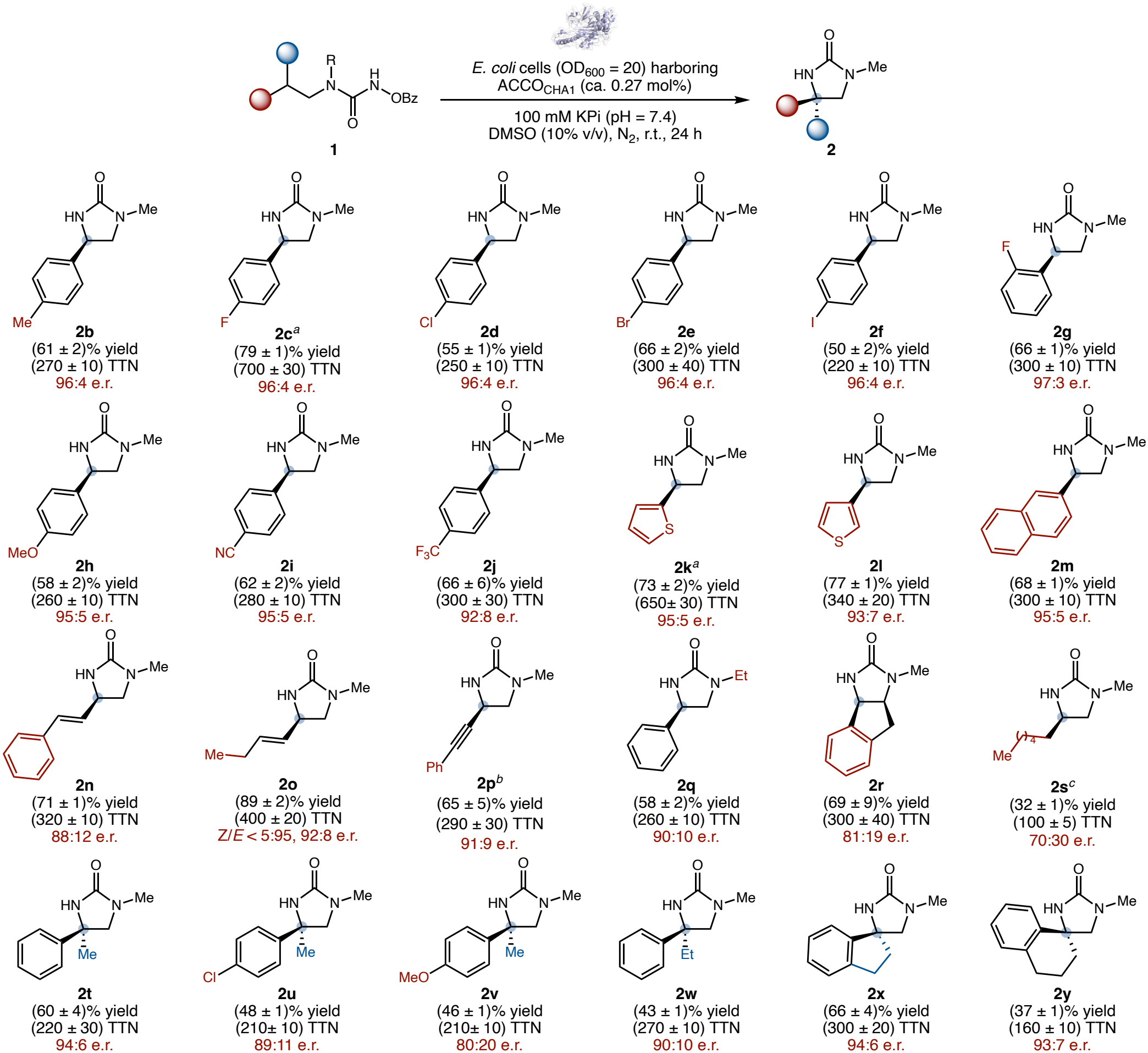
Substrate Scope of ACCO_CHA1_-Catalyzed Enantioselective C(sp^3^)–H Amidation. Reaction conditions: **1** (10.0 mM), 360 μL *E. coli* cells harboring ACCO_CHA1_ (OD_600_ = 20) in KPi buffer (100 mM, pH = 7.4, ca. 0.27 mol% ACCO_CHA1_), DMSO (10% v/v), room temperature, 24 h. All the biocatalytic reactions were performed as technical triplicates, *n* = 3; reported yield data represent mean ± s.d. of triplicate yields. TTN calculation was based on enzyme concentrations in the cell-free lysates determined by SDS–PAGE analysis in triplicate with an internal standard, *n* = 3; reported TTN data represent (mean ± s.d.) of triplicate yields and triplicate enzyme concentrations. See the SI for detailed experimental procedures. ^*a*^Using *E. coli* cells harboring ACCO variants (OD_600_ = 10) in KPi buffer (100 mM, pH = 7.4, ca. 0.13 mol% ACCO_CHA1_). ^*b*^Using ACCO_Nim2_ L250F in lieu of ACCO_CHA1_. ^*c*^Using *E. coli* cells harboring ACCO _CHA1_ (OD_600_ = 30) in KPi buffer (100 mM, pH = 7.4, ca. 0.39 mol% ACCO_CHA1_).

With results of secondary C(sp^3^)–H amidation in hand, we next investigated the use of ACCO_CHA1_ for the biocatalytic enantioconvergent tertiary C(sp^3^)–H amidation using racemic substrates. Gratifyingly, without additional enzyme engineering, ACCO_CHA1_ effected the conversion of (*rac*)-**1t** into **2t** with excellent enantioselectivity (60% yield, 94:6 e.r.). Both electron-withdrawing substituents such as a *para*-chloro group (**2u**) and electron-donating substituents such as a *para*-methoxy group (**2v**) on the aromatic ring were tolerated. The C4-ethyl (**2w**) congener was also converted by ACCO_CHA1_, although with slightly reduced yield and enantioselectivity. Furthermore, nonheme Fe biocatalyst ACCO_CHA1_ enabled the enantioconvergent construction of spirocyclic molecular architectures. Substrates derived from indane (**1x**) and tetralin (**1y**) were effectively transformed into the corresponding 5,5- and 5,6-spirocyclic products **2x** and **2y**, respectively. Collectively, the ability of ACCO_CHA1_ to functionalize several types of secondary and tertiary C(sp^3^)–H bonds indicated its synthetic potential as a versatile biocatalyst for constructing structurally diverse enantioenriched 1,2-diamine derivatives using a C–H functionalization logic.

Having developed ACCO_CHA1_ for the asymmetric synthesis of a wide range of 1,2-diamine derivatives, we next sought to engineer nonheme Fe enzymes for the regiodivergent enantioselective synthesis of both 1,2- and 1,3-diamines from common precursors via C(sp^3^)–H amidation in a biocatalyst-controlled fashion. To this end, **4a**, possessing both unactivated and benzylic C(sp^3^)–H bonds, was selected as the model substrate. Starting from **4a**, C(sp^3^)–H amidation through a 1,5-HAT or a 1,6-HAT pathway would furnish 2-imidazolidinone **5a** or 2-tetrahydropyrimidinone **6a**, respectively. Our DFT calculated C(sp^3^)–H bond dissociation energies indicated an intrinsic difference of 11.4 kcal/mol, indicating potential challenges in achieving selective transformations. The use of ACCO_CHA1_ for the C(sp^3^)–H amidation of **4a** afforded **5a** in a modest 28% yield with 77:23 e.r., along with **6a** in 3% yield and 92:8 e.r. This initial result indicated the feasibility of achieving biocatalyst-controlled regiodivergence. Screening our in-house collection of *Phy*ACCO variants identified ACCO_Nim2_ and ACCO I184A K158T K172V L186V as the most selective starting points for the formation of imidazolidinone **5a** and tetrahydropyrimidinone **6a**, respectively. In particular, ACCO_Nim2_ favored **5a** (63% yield and 78:22 e.r.) with a 90:10 ratio of **5a**:**6a**. The formation of the 1,3-diamine derivative **6a** was found to be more challenging, and ACCO I184A K158T K172V L186V was found to catalyze the formation of **6a** in 11% yield and 88:12 e.r., albeit with a **5a**:**6a** ratio of 78:22 still favoring **5a** (see Table S8 for details). Encouraged by these results, we sought to develop site-selective biocatalysts for regiodivergent C(sp^3^)–H amidation through directed evolution.

Analysis of ACCO lineage screening results suggested that residue 158, located on a β-strand proximal to the nonheme Fe center, plays a critical role in controlling both site- and enantioselectivity (Figure 4B). We therefore performed SSM and screening by targeting T158 using ACCO_Nim2_ as the parent. Beneficial mutation T158W was found to enhance the C(sp^3^)–H amidation regioselectivity for **5a**, increasing the **5a**:**6a** ratio to 96:4 while also improving the enantioselectivity of **5a** from 78:22 e.r. to 84:16 e.r.. In subsequent rounds of directed evolution, additional residues on β-strands surrounding the Fe center were targeted, resulting in the identification of S160M and V186C as beneficial substitutions that further improved the enantioselectivity of **5a**. Based on our previous observation that more distal residues lining the substrate-access tunnel can also influence catalytic performance, we performed SSM and screening by randomizing F91 at the entrance of the substrate tunnel distal to the Fe center (Cα···Fe distance = 15.0 å). In this final round of engineering, F91L emerged as a beneficial mutation, increasing the combined C(sp^3^)–H amidation yield to (75 ± 2)%, with a **5a**:**6a** ratio of 93:7 and a 95:5 e.r. for the major product **5a** (Figure 4A, left; see Table S9 for details). This evolved variant was designated as ACCO_CHA2_ (ACCO C–H amidase 2).

**Figure 4.**
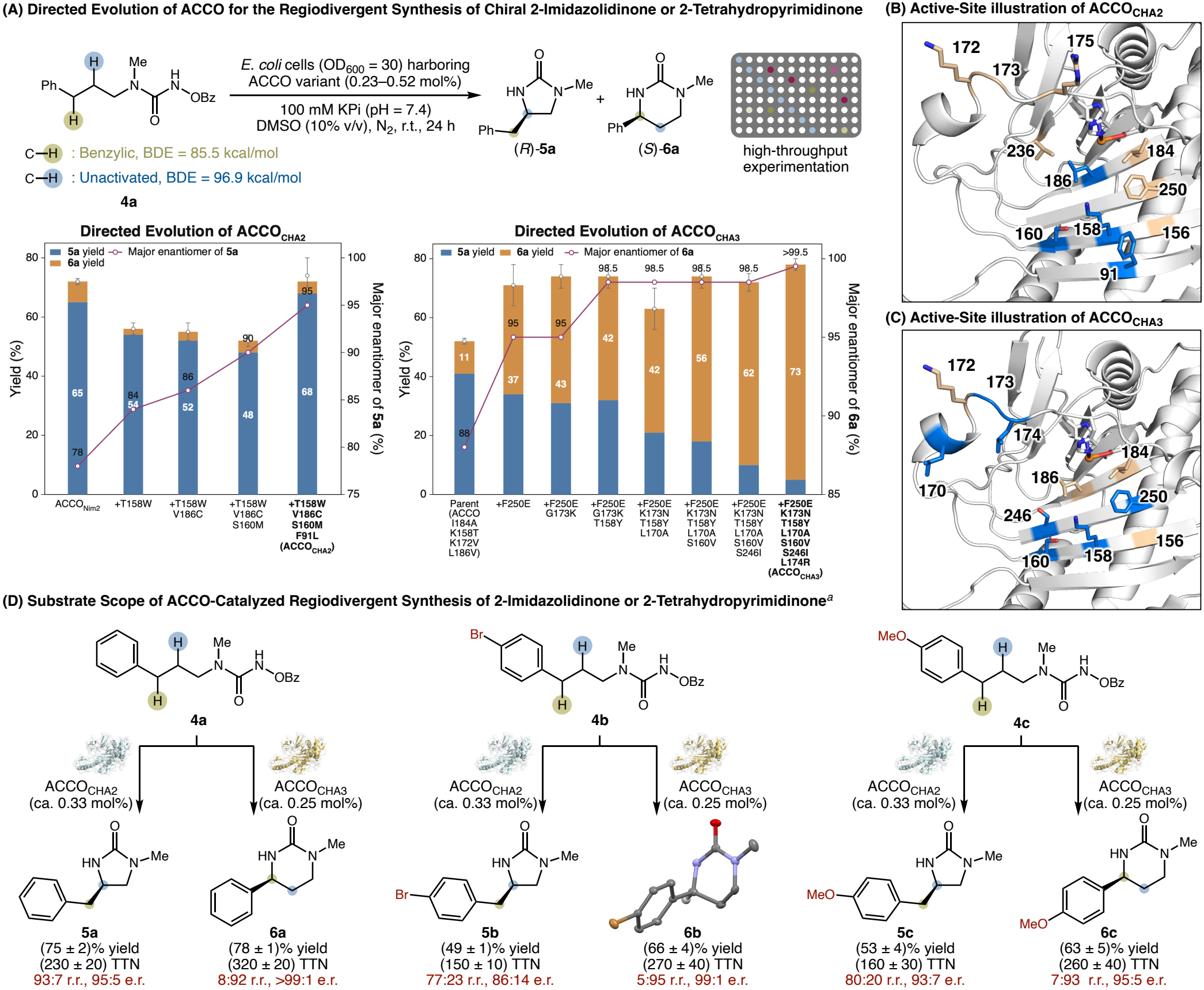
Directed Evolution of ACCO_CHA2_ and ACCO_CHA3_ for the Asymmetric Synthesis of α 2-Imidazolidinone or 2-Tetrahydropyrimidinone via Selective C(sp^3^)–H Functionalization. **(A)** Directed evolution of ACCO for the stereodivergent synthesis of chiral 2-imidazolidinone or 2-tetrahydropyrimidinone using whole *E. coli* cells. Bond dissociation energies (BDEs) shown above were calculated at the M06−2X/def2-TZVP level of theory, see the SI for computational details. **(B)** Active-site illustration of ACCO (PDB ID: 1WA6) and beneficial mutations identified from directed evolution of ACCO_CHA2_. Marine residues denote beneficial mutations identified in ACCO_CHA2_. Wheat residues denote mutations of ACCO_Nim2_ relative to wt ACCO. **(C)** Active-site illustration of ACCO (PDB ID: 1WA6) and beneficial mutations identified from directed evolution of ACCO_CHA3_. Marine residues denote beneficial mutations identified in ACCO_CHA3_. Wheat residues denote mutations of ACCO I184A K158T K172V. **(D)** Substrate scope of the synthesis of 2-imidazolidinone or 2-tetrahydropyrimidinone via selective amidation of activated or unactivated C(sp^3^)–H bonds. Reaction conditions: 1 (10.0 mM), 360 μL *E. coli* cells harboring ACCO variants in KPi buffer (OD_600_ = 30, 100 mM, pH = 7.4), DMSO (10% v/v), room temperature, 24 h. All the biocatalytic reactions were performed as technical triplicates, *n* = 3; reported yield data represent mean ± s.d. of triplicate yields. TTN calculation was based on enzyme concentrations in the cell-free lysates determined by SDS–PAGE analysis in triplicate, *n* = 3; reported TTN data represent mean ± s.d. of triplicate yields and triplicate enzyme concentrations. See the SI for the detailed experimental procedure.

In parallel, ACCO I184A K158T K172V L186V was selected as the parent for evolving a biocatalyst toward the regioselective asymmetric synthesis of 1,3-diamine derivative **6a**. Similar to the engineering of ACCO_CHA2_, SSM and screening targeting active-site and substrate tunnel residues were used as the key engineering strategy. The first round of engineering identified F250E as a key beneficial substitution, substantially improving site selectivity, enantioselectivity, and enzyme activity. The resulting variant ACCO I184A K158T K172V L186V F250E afforded **5a** and **6a** in a 71% combined yield with a 48:52 ratio of **5a**:**6a** and a 95:5 e.r. for **6a**. Two additional rounds of directed evolution identified G173K and T158Y as beneficial substitutions, increasing the yield and e.r. of **6a** to 57% and 98.5:1.5. During the next round of engineering randomizing L170, we identified a variant containing both L170A and an unanticipated mutation, K173N. Although G173N alone was not identified as a beneficial mutation in the preceding round, its combination with L170A improved the **5a**:**6a** ratio to 34:66, while maintaining a 98.5:1.5 e.r. for **6a**. Further SSM and screening identified S160V and S246I, both located on β-strands close to the Fe center, as additional beneficial substitutions. Replacing the corresponding serine with hydrophobic residues valine and isoleucine at these two sites further enhanced selectivity for **6a**, affording a 14:86 ratio of **5a**:**6a** in 72% combined yield while retaining a 98.5:1.5 e.r. for **6a**. Finally, incorporation of a beneficial mutation L174R from the 172–182 loop further improved both site- and enantioselectivity, affording the final variant ACCO I184A K172V L186V F250E G173N K158Y L170A S160V S246I L174R, designated ACCO_CHA3_. At a loading of 0.25 mol%, ACCO_CHA3_ furnished 2-tetrahydropyrimidinone **6a** in (73 ± 2)% yield with a **5a**:**6a** ratio of 7:93 and >99:1 e.r. of **6a**. Together, the successful engineering of ACCO_CHA2_ and ACCO_CHA3_ demonstrated the power of directed evolution to override the innate substrate control over reaction site selectivity in asymmetric C(sp^3^)–H amidation reactions.

Having established biocatalytic control over 1,2- and 1,3-diamine derivative synthesis, we next evaluated the substrate scope of regiocomplementary biocatalysts ACCO_CHA2_ and ACCO_CHA3_ (Figure 4(D)). For substrate **4b** bearing a *para*-bromophenyl group, ACCO_CHA2_ selectively furnished the 1,2-diamine product **5b** in (49 ± 1)% yield, 77:23 regioisomeric ratio (r.r.), and 86:14 e.r.. ACCO_CHA3_ also catalyzed the synthesis of the 1,3-diamine product **6b** in (66 ± 4)% yield, 5:95 r.r., and 99:1 e.r.. Substrate **4c** possessing an electron-donating *para*-methoxyphenyl group also underwent regiocomplementary C(sp^3^)–H amidation with enzyme-controlled site selectivity. The use of ACCO_CHA2_ afforded **5c** in (53 ± 4)% yield, 80:20 r.r., and 93:7 e.r., while the use of ACCO_CHA3_ led to **6c** in (63 ± 5)% yield, 7:93 r.r., and 95:5 e.r.. Finally, the absolute stereochemistry of 1,3-diamine product **6b** was determined by single crystal X-ray diffraction analysis.

To better understand the catalytic efficiency of ACCO_CHA1_, we determined its total turnover numbers (TTNs) using purified enzyme at progressively decreased catalyst loadings (Figure 5A). At an enzyme loading of 0.30 mol%, ACCO_CHA1_ furnished (*R*)-**2a** in (79 ± 2)% yield and 97:3 e.r., corresponding to a TTN of 260 ± 7. Decreasing the enzyme loading to 0.10 and 0.025 mol% afforded **2a** in (69 ± 6)% and (55.2 ± 0.1)% yield, respectively, while maintaining a 97:3 e.r. These conditions corresponded to TTNs of 690 ± 60 and 2,210 ± 10, respectively. Importantly, ACCO_CHA1_ remained catalytically competent at a loading of 0.010 mol%, providing **2a** in (45 ± 6)% yield, 96:4 e.r. and (4,500 ± 600) TTN. Further decreasing enzyme loading did not lead to an increase in TTN (Table S17). Together, these results established a TTN of ca. 4,500 for engineered nonheme biocatalyst ACCO_CHA1_. This total turnover number is approximately 90-fold higher than a previously developed, well-tailored small-molecule Fe catalysts^9g^ for this type of C(sp^3^)–H amidation reaction, demonstrating the catalytic efficiency achievable with nonheme Fe enzymes.

**Figure 5.**
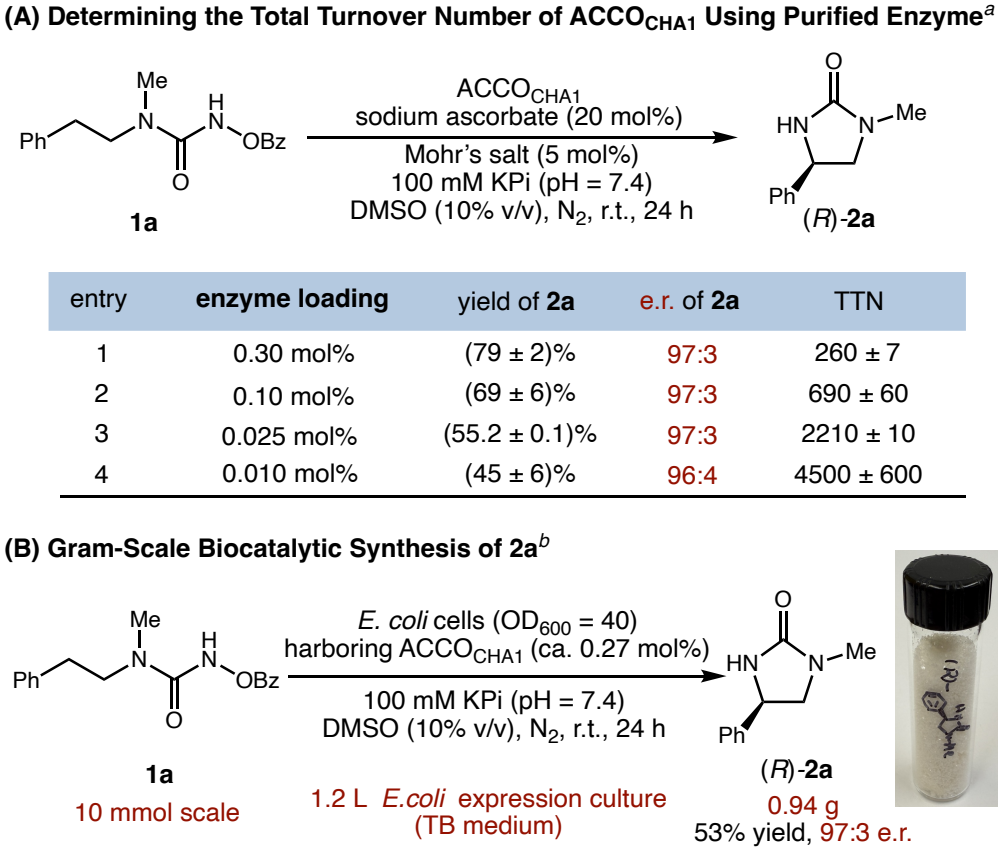
Evaluating the Total Turnover Numbers (TTNs) and Gram-scale Biotransformations of ACCO_CHA1_ in Biocatalytic Synthesis of 2-Imidazolidinone (2a). ^*a*^Reaction conditions: 10 mM **1a**, 0.5 mM (NH_4_)_2_Fe(SO_4_)_2_, 2 mM sodium ascorbate, 360 μL 100 mM KPi buffer (pH = 7.4), enzyme loadings are as depicted in the table. All biocatalytic reactions were performed as technical triplicates, *n* = 3; reported yield data represent mean ± s.d. of triplicate yields. ^*b*^Reaction conditions: 10.0 mmol **1a**, 500 mL *E. coli* cells harboring ACCO_CHA1_ in KPi buffer (OD_600_ = 40, 100 mM, pH = 7.4), DMSO (10% v/v), room temperature, 24 h.

To further demonstrate the synthetic utility of this biocatalytic C(sp^3^)–H amidation, we carried out this biotransformation on a 10.0 mmol scale using whole *E. coli* cells overexpressing ACCO_CHA1_ (Figure 5(B)). The cell pellet obtained from a 1.2 L expression TB culture was employed directly in this 10.0 mmol-scale biotransformation. Under the standard reaction conditions, 0.94 g of (*R*)-**2a** was isolated in 53% yield with 97:3 e.r., demonstrating the scalability of the current whole-cell biotransformation.

To explore the origin of the regioselectivity controlled by enzymes, we performed density functional theory (DFT) and classical molecular dynamics (MD) simulations to investigate active site residue-substrate interactions (see SI for details). These results suggest that 1,6-HAT is electronically favored owing to the weaker benzylic C(sp^3^)–H bond, which leads to a stabilized benzylic radical intermediate. This preference is further reinforced by a cation-π interaction between the phenyl group of the substrate and R174 residue in ACCO_CHA3_. In contrast, in ACCO_CHA2_, a distinct π-π interaction between the substrate’s phenyl group and W158, together with an additional C–H…π interaction with M160, locks the phenyl group in a reactive conformation that favors 1,5-HAT, affording the 1,2-diamine product.

To further investigate the mechanism of this nonheme Fe enzyme-catalyzed C(sp^3^)–H amidation, we performed a series of experiments (Figure 6). First, addition of the radical scavenger TEMPO inhibited product formation in a concentration-dependent manner. Whereas **2a** was obtained in (79 ± 1)% yield in the absence of TEMPO, its yield decreased to (53 ± 6)% upon the addition of 1.0 equiv of TEMPO and to (8 ± 2)% upon the addition of 2.0 equiv of TEMPO (Figure 6(A)). In addition, TEMPO-trapping adduct **8** was detected by LC–MS. These results indicated the involvement of a carbon-centered radical intermediate in this nonheme Fe enzyme-catalyzed C(sp^3^)–H amidation.

**Figure 6.**
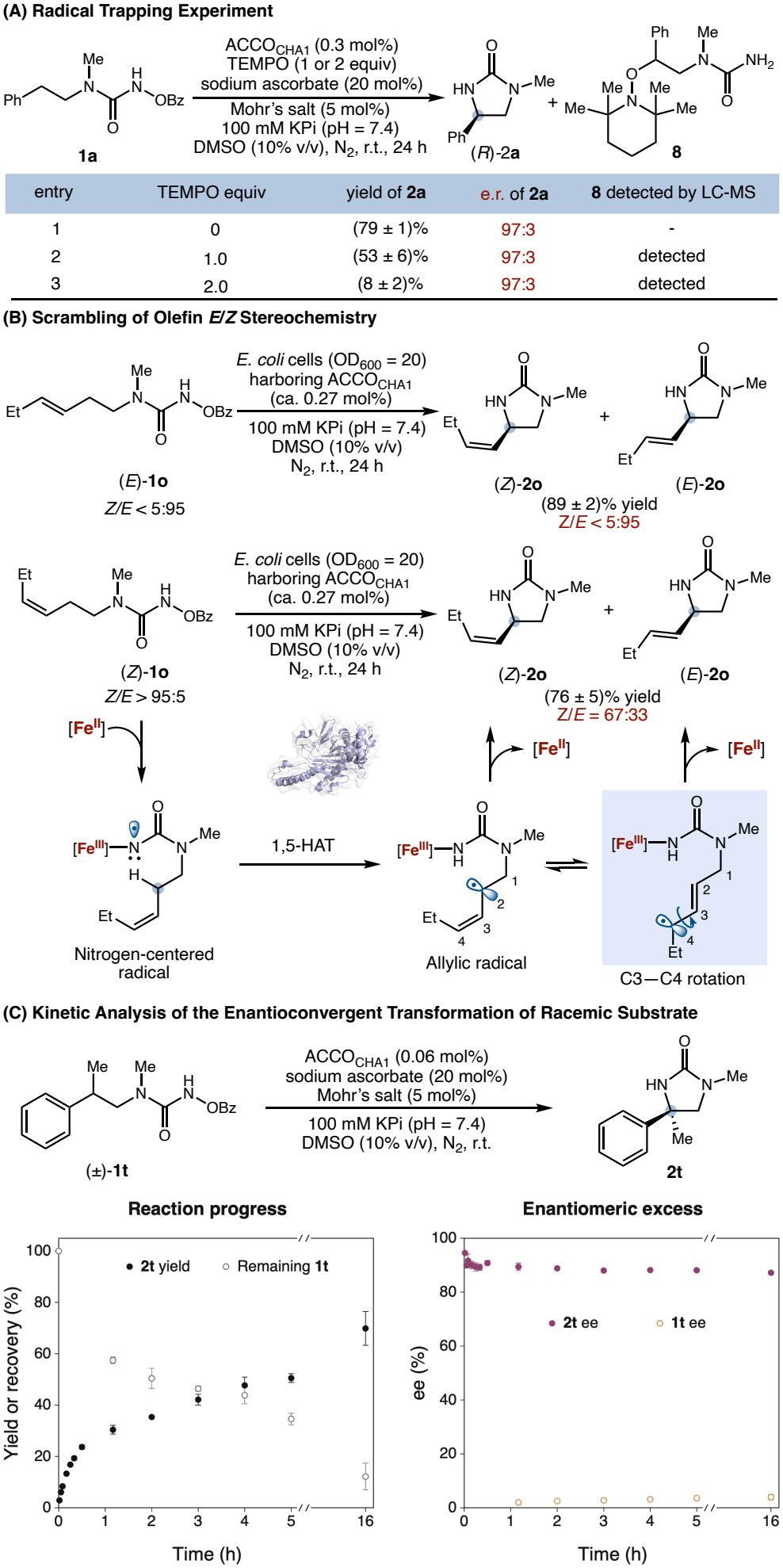
Mechanistic Studies Indicative of a Radical-Mediated Mechanism. (A) TEMPO trapping experiment. (B) Scrambling of olefin stereochemistry during biocatalytic C(sp^3^)–H amidation. (C) Biocatalytic transformation of racemic tertiary C(sp^3^)–H substrate **1t.** Triplicated results are shown (see the SI for details).

To further probe the mechanism of this nonheme Fe enzyme-catalyzed amidation, we prepared geometrically well-defined allylic substrates (*E*)- and (*Z*)-**1o** and subjected them to the biocatalytic conditions (Figure 6(B)). Biotransformation of (*E*)-**1o** (*E*/*Z* > 95:5) afforded the corresponding allylic C–H amidation product **2o** with complete retention of olefin configuration (*E*/*Z* > 95:5). In contrast, the use of geometrically pure (*Z*)-**1o** (*Z*/*E* > 95:5) furnished **2o** with a *Z*/*E* ratio of 67:33, indicating scrambling of olefin stereochemistry during this biocatalytic C(sp^3^)–H amidation process. The partial erosion of olefin stereochemical integrity is consistent with a stepwise, radical-mediated mechanism. Previous studies showed that Fe nitrenoid can be viewed as a diradical species, with substantial radical character on the nitrogen center.^20–22^ Hydrogen atom transfer between the enzymatic Fe nitrenoid and the allylic C(sp^3^)–H substrate generates an allylic carbon-centered radical and an Fe(III)–NHR moiety. The delocalized nature of the allylic radical allows rotation along the C3–C4 axis, leading to the scrambling of olefin geometry prior to the C–N bond-forming radical rebound. Through this radical mechanism, the less thermodynamically stable (*Z*)-**1o** was partially converted to the thermodynamically more stable (*E*)-**2o** product. Importantly, the partial retention of the starting (*Z*)-configuration indicates that the radical rebound step effectively competes with allylic σ–π–σ isomerization and is therefore comparatively rapid within the ACCO_CHA_ active site. This result is distinct from the previously reported Fe complex-^9f,g^ or heme-dependent enzyme-^4d,m,r^ catalyzed C(sp^3^)–H amidation, in which a (*Z*)-configured substrate furnished exclusively the corresponding *E*-configured products. The incomplete *Z*-to-*E* isomerization with ACCO_CHA_ indicated a shorter-lived carbon-centered radical in ACCO_CHA_ compared to those with small-molecule Fe complex and myoglobin.

With experimental evidence in support of a radical-mediated mechanism, we further investigated the stereochemical course of the biocatalytic transformation of the racemic tertiary C(sp^3^)–H substrate (±)-**1t** (Figure 6(C)). Our studies showed that the formation of the C–H amidation product **2t** was accompanied by progressive consumption of the substrate **1t**. Throughout the course of this biotransformation, the unreacted **1t** remained nearly racemic, with only a minor change in its enantiomeric ratio from 50:50 to 52:48. Similarly, product **2t** formed with consistently high enantioselectivity, showing only a slight decrease in enantioenrichment from 95:5 e.r. (3 min) to 94:6 e.r. (16 h). These results established the enantioconvergent nature of this transformation for tertiary C(sp^3^)–H substrates.

To determine the kinetic significance of the C–H cleavage step, we next determined the kinetic isotope effect (KIE) by independently measuring the initial rates of biocatalytic C–H amidation using **1a** and its doubly deuterated analog **1a**-*d*_2_ (Figure 7(A)). These initial rate measurements revealed a primary KIE (*k*_H_/*k*_D_) of 3.5 ± 0.2, indicating that C–H cleavage is the rate-determining step.

**Figure 7.**
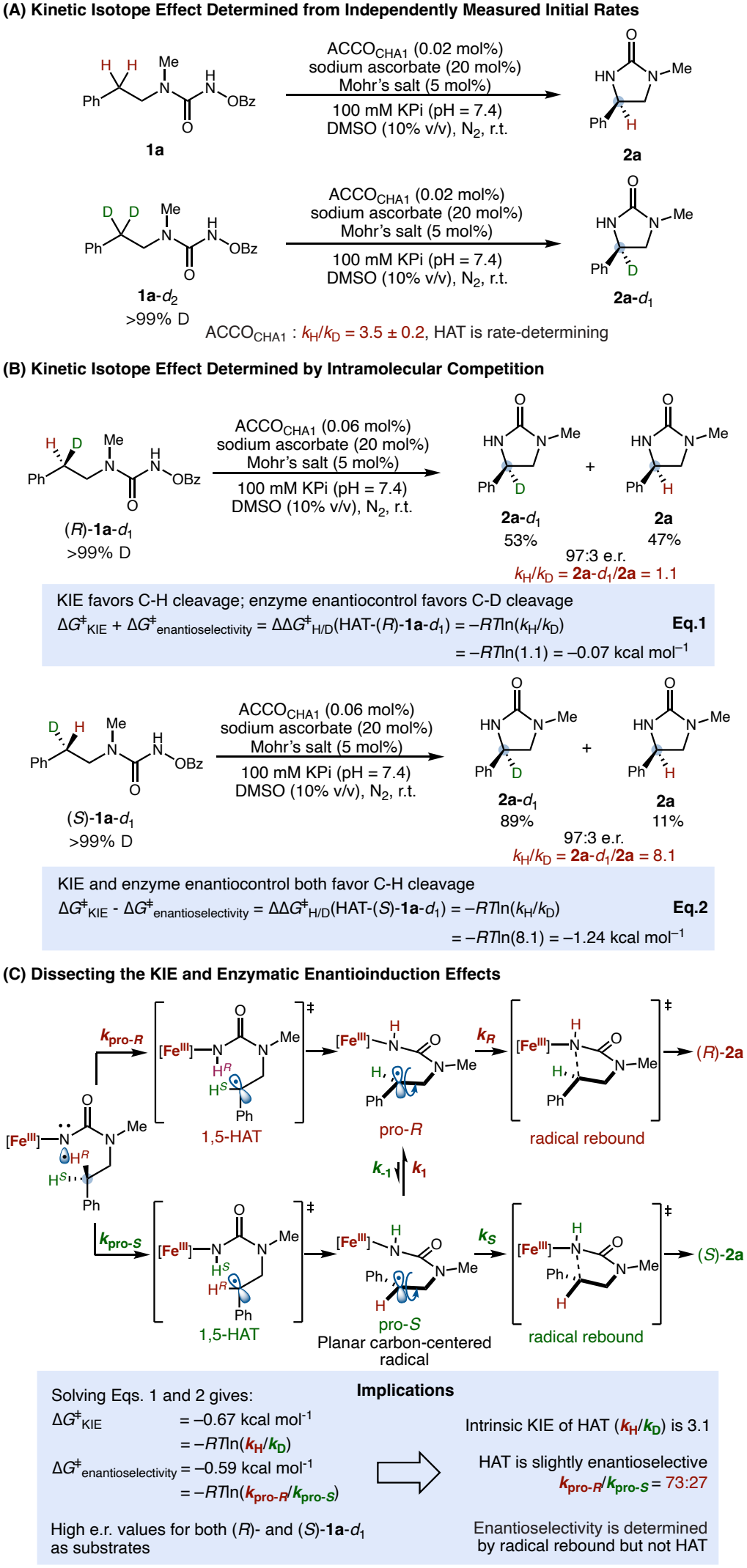
Kinetic Isotope Effect Studies. Reaction conditions: 10 mM 1a (or isotopically labeled analogs), 0.5 mM (NH_4_)_2_Fe(SO_4_)_2_, 2 mM sodium ascorbate, 0.02 mol% or 0.06 mol% purified ACCO_CHA1_, 100 mM KPi buffer (pH = 7.4). See the SI for a detailed experimental procedure.

To gain further insights into the rate-limiting step and the enzymatic stereoinduction during the hydrogen atom transfer step, we prepared the two stereoisotopomers (*R*)-**1a**-*d*_1_ and (*S*)-**1a**-*d*_1_ and subjected them to the biocatalytic reaction conditions (Figure 7(B)). Previous studies on related nonheme^11a,e^ and heme^22^ Fe enzyme systems showed that the HAT step is both exergonic and irreversible. Thus, the contributions of the intrinsic kinetic isotope effect (KIE) and enzymatic enantiopreference during the HAT step in this intramolecular competition could be dissected for both (*R*)-**1a**-*d*_1_ and (*S*)-**1a**-*d*_1_ by carefully measuring the distribution of **2a**-*d*_1_ and **2a** (Figure 7(B) and 7(C)).

For (*R*)-**1a**-*d*_1_, the intrinsic preference for C–H cleavage opposed the enzymatic enantiopreference, which favored cleavage of the competing C–D bond. Consequently, the two products formed in a nearly equal ratio of 53:47 for **2a**-*d*_1_:**2a** based on ^1^H NMR spectroscopic analysis, corresponding to an apparent *k*_H_/*k*_D_ of 1.1 and an activation energy difference of −0.07 kcal mol^-1^. In contrast, for (*S*)-**1a**-*d*_1_, the KIE and enzymatic enantiopreference both favored C–H cleavage. These reinforcing effects afforded a **2a**-*d*_1_:**2a** ratio of 89:11, corresponding to an apparent *k*_H_/*k*_D_ of 8.1 and an activation energy difference of −1.24 kcal mol^-1^. Dissection of these opposing and reinforcing effects gave free-energy contributions of approximately −0.67 kcal mol^-1^ for the intrinsic kinetic isotope effect (Δ*G*^‡^ _KIE_) and −0.59 kcal mol^-1^ for the enzymatic enantiopreference during HAT (Δ*G*^‡^ _enantioselectivity_). The latter corresponds to a modest HAT enantioselectivity of approximately 73:27 (*k*_pro-*R*_/*k*_pro-*S*_). Nevertheless, both stereoisotopomers furnished (*R*)-**2a** with similarly high enantioselectivity (97:3 e.r.), demonstrating that the product configuration is not determined during the HAT step. Instead, formation of a planar carbon-centered radical and the facile interconversion between the pro-(*R*) and pro-(*S*) states of the radical partially ablates the stereochemistry at the reacting carbon. Ultimately, the overall reaction enantioselectivity is determined by the highly stereoselective C−N bond-forming radical rebound step. Collectively, these results suggest the HAT step as the rate-determining step and the radical rebound step as the enantioselectivity-determining step for this nonheme Fe enzyme-catalyzed C(sp^3^)–H amidation reaction.

## Conclusion

In summary, we have repurposed and evolved the nonheme Fe enzyme ACCO into a family of highly selective C(sp^3^)–H amidases for the asymmetric synthesis of chiral 1,2- and 1,3-diamine derivatives. Directed evolution furnished complementary biocatalysts capable of catalyst-controlled, regiodivergent C–H amidation through selective 1,5- and 1,6-hydrogen atom transfer, overcoming a challenge in site-selective C–H functionalization. The engineered enzymes exhibited a broad substrate scope, enabled enantioconvergent functionalization of tertiary C(sp^3^)–H bonds, and achieved total turnover numbers of up to 4,500. Mechanistic studies indicated a stepwise radical pathway, in which C–H bond cleavage constitutes the rate-limiting step, whereas the stereochemistry is established during the subsequent C–N bond-forming radical rebound step. More broadly, this work established nonheme Fe enzymes as a versatile platform for nitrene transfer catalysis beyond recently developed nitrogen migration reactions and demonstrated that directed evolution could simultaneously control reaction site selectivity and stereoselectivity in radical-mediated C–H functionalization, opening new opportunities to develop selective nonheme enzymes for non-native biocatalysis.

## Acknowledgements

We are grateful to the National Institutes of Health (R35GM147387 for metalloenzyme research) and the National Science Foundation (CHE-2400087 for mechanistic study) for financial support. Y.Y. is an Alfred P. Sloan Research Fellow (FG-2024-22244), a Camille Dreyfus Teacher-Scholar Awardee (TC-25-084), a David & Lucile Packard Fellow (2023-76169) and a

Howard Hughes Medical Institute Freeman Hrabowski Scholar. Computations were carried out at the University of Pittsburgh Center for Research Computing and Data and the Advanced Cyberinfrastructure Coordination Ecosystem: Services & Support (ACCESS) Program, supported by NSF grants OAC-2138259, OAC-2138286, OAC-2138307, OAC-2137603, OAC-2138296 and CHE-260005.

## Notes

### Competing Interest Statement

The authors have declared no competing interest.

### Summary of Updates

We revised the manuscript to improve clarity and readability. We also updated the figures.

